# Long time-scale study of von Willebrand factor multimers in extensional flow

**DOI:** 10.1101/2020.09.09.290304

**Authors:** S. Kania, A. Oztekin, X. Cheng, X. F. Zhang, E. B. Webb

**Affiliations:** Lehigh University

**Keywords:** Weighted Ensemble Brownian dynamics, rare events, non-equilibrium system, vWF, extensional flow

## Abstract

Extensional flow-induced transitions from a compact to an unfolded conformation are explored for the human glycoprotein von Willebrand factor (vWF). Multimer unfolding is a crucial step in the process of blood clotting and protein size maintenance. Previous studies have shown that flow-induced conformational transitions are initiated by a thermally nucleated polymeric protrusion. Below a certain strain rate, such a transition is a rare event that cannot be studied using standard stochastic dynamic simulation. In the present study, we have employed Weighted Ensemble Brownian dynamic (WEBD) simulations to study rare events of conformation transition in extensional flow. Results are presented for the transition rate of VWF multimer unfolding, with concomitant analysis of the likelihood of pathological unfolding as a function of strain rate. Relative to the typical half-life of vWF proteins in the human body, results here indicate that pathological unfolding would not manifest for strain rate less than 2000 s^−1^.

**Statement of Significance:** vWF multimers, as they transit through the circulation, are exposed to extensional flow multiple times, and the total exposure time to such intermittent extensional flow can be on the order of minutes to an hour. However, due to the time-scale limitation of Brownian dynamics simulation, all the present studies of vWF multimers are limited to a few seconds in total duration. Here, we have applied an enhanced sampling technique, i.e., Weighted Ensemble, in combination with Brownian dynamics to analyze the behavior of multimers in extensional flow at physiologically relevant time-scales of hours and longer. The findings presented here provide new physical insights into vWF behavior, including how it relates to hematological pathology, while also illustrating the time-scale bridging capability of the WEBD method.

## Introduction

When blood vessels are injured, there is a drastic change in the blood flow rate. One such severe result is excessive bleeding, which can be lethal if blood clot formation is unable to occur. However, the increase in blood flow rate helps to initiate clot formation with the help of the blood protein, von Willebrand factor (vWF)(1, 2). vWF senses the variation in the blood flow at the site of vascular injury, and it unravels from globular to elongated conformation(3). Once stretched, vWF is capable of adhering to the surface of a damaged blood vessel via a binding mechanism to collagen while simultaneously binding to platelets in the blood(4, 5). In this way, vWF initiates the formation of a hemostatic plug. This extraordinary response of vWF to changes in blood flow is critically dependent on the protein’s multimeric size (i.e., the number of repeat units that comprise a vWF protein)(6). Hence, the abnormal size distribution of vWF in blood plasma can lead to various diseases.

vWF is secreted into blood plasma as ultra-large vWF (i.e., ~1000 monomers), which are unstable. In normal condition, this ultra-large vWF is cleaved by ADAMTS13 enzyme to achieve the functional size distribution (~ 40-200 monomers) of vWF multimers in blood plasma(7, 8). Absence of ADAMTS13 results in a high concentration of large vWF leading to a life-threatening disease caused by uncontrolled microvascular thrombosis(9, 10). On the other hand, excessive cleavage by ADAMTS13 results in a lack of multimer in functional size range, resulting in bleeding disorder knowns as type 2A von Willebrand disease(11). Thus, cleavage by ADAMTS13 of vWF is a vital regulatory mechanism. ADAMTS13 cleaves vWF by breaking the Tyr1605-Met1606 bond in one of the protein’s A2-domains(8, 11, 12). Similar to the vWF proteins, or multimers, the A2 domain in each monomer is in a compact folded conformation unless subject to sufficient tensile force. The bond at which ADAMTS13 cleaves vWF, is deeply buried in the folded A2-domain(13, 14). For wild type vWF, in the absence of applied tensile loading, A2-domain unfolding is a rare event; however, the likelihood of unfolding is directly related to the tensile force exerted on the domain(15). Moreover, sufficient tensile force can only be exerted on A2 domains after the vWF multimer unravels to an elongated conformation(16). Hence, the unraveling of vWF multimers is not only crucial for the hemostasis process, it is also essential for scission by ADAMTS13. Herein, the conformational change from compact, or folded, to elongated, or unfolded, will be called unfolding for A2 domains and unraveling for vWF protein multimers. To be clear, the unraveled polymer state is when the multimer becomes completely elongated along the flow direction.

In order to elucidate the unraveling process of vWF in different flow conditions, many authors performed Brownian dynamic (BD) simulations of vWF as a collapsed polymer via a bead-spring model(5, 17, 18). Alexander-Katz et al. have explained the flow caused conformational changes (globular to unraveled) of vWF by a nucleation model based on the presence of thermally nucleated polymeric protrusions(19,20). Protrusion nucleation theory was further verified by the presence of a double-peak structure in an averaged tensile force profile for multimers subjected to shear flow(21). Figure 1 illustrates a globular to unraveled transition observed in BD simulations presented here; also shown are examples of thermal protrusion of globular state polymer, similar to the one that eventually nucleates the transition. When a thermal protrusion extends far enough from the globule to be pulled by the surrounding flow field, it results in a half-dumbbell geometry, which is very unstable state (i.e., spontaneous unraveling) as shown in Fig. 1.

**Figure 1:**
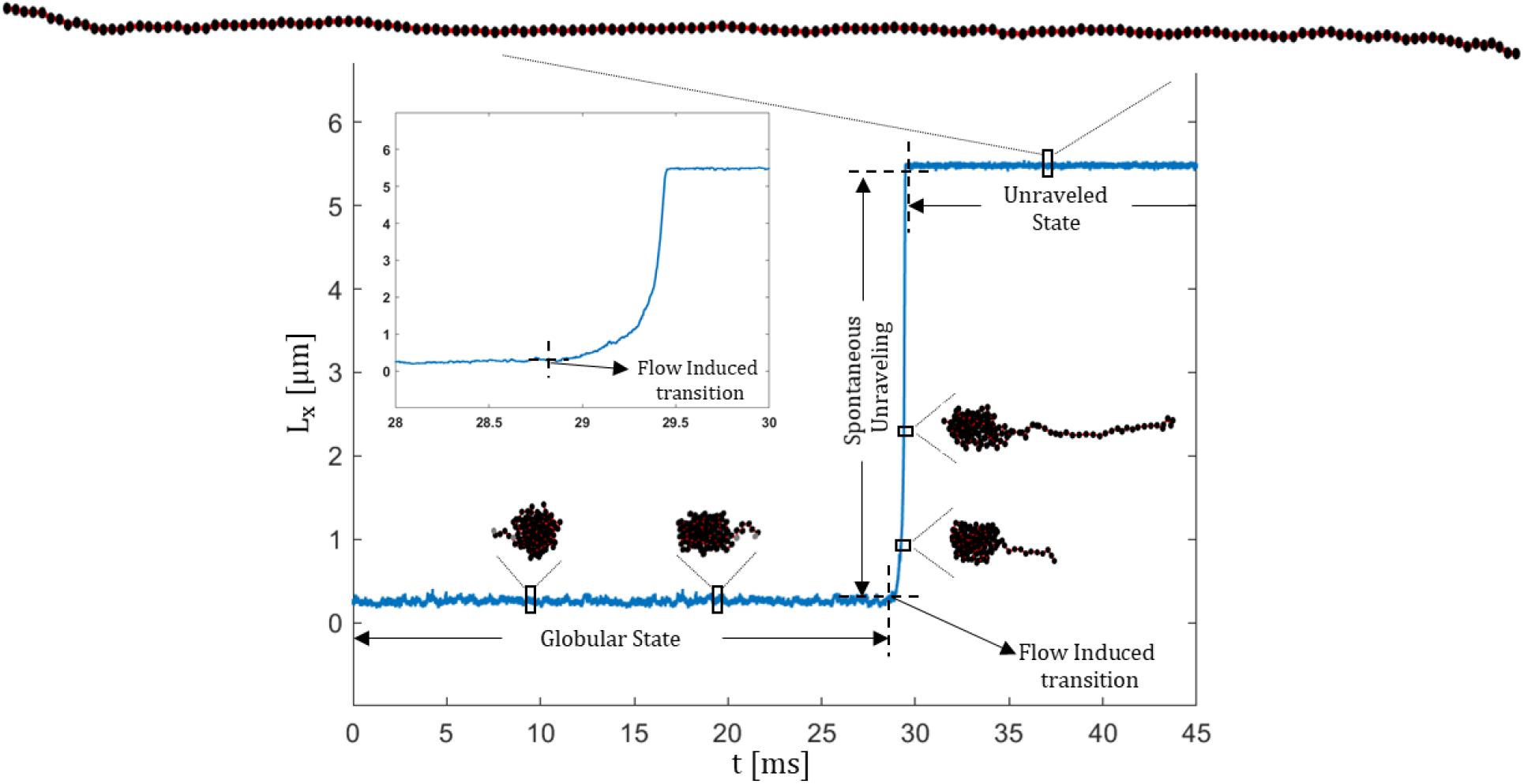
Plot of polymer extension (L_x_) in flow direction as a function of time from Brownian dynamic simulation at the extensional rate 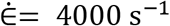. The time duration for which collapsed polymer is in the globular state and the time where flow caused spontaneous unraveling are indicated by arrows. The inset represents the time duration of spontaneous unravelling. The small rectangles in the varied states (globular, spontaneous unraveling, unraveled state) denote the configurations from which the snapshots were taken. The snapshots of collapsed polymer (globular state) indicates the start and the end of protrusion with gray-colored bead.

Prior work of our group indicated that extensional flow is considerably more effective in unraveling the multimer than pure shear flow(22). It further gave evidence that vWF multimers within the functional size range unravel from globular conformation for elongation rates higher than 3500 s^−1^ (i.e., such as what would be encountered in the vicinity of vascular injury)(22). For extensional rates below the identified transition rate, the transition from globular to unravel state is a rare event whose likelihood decreases with decreasing extension rate.

Despite much excellent work in understanding the unraveling process of vWF molecules in varying flow conditions, research has not yet fully explored unraveling of vWF multimers at low extensional rates where conformation changes are rare events with regard to BD simulation time scales. While in principle, it is possible to run BD simulation for any duration of time, in practice this is not possible due to the limitation of computational resources. Couple to this is the physiological time scale of vWF protein in human blood stream; the half-life of functional size vWF in blood is reported to be 12 hours(23, 24). As such it is possible for vWF protein to exist in the blood stream for order of 1-2 days. Therefore, condition exist that causes functional size vWF to unravel and scission on time scale shorter than its life time i.e., minutes/hours, and this can be the source of pathology. Unfortunately, employing BD simulation to sample event on order of minutes or even few seconds becomes prohibitive. This is the case for relatively lower extensional rate; where healthy size vWF may undergo unraveling and scission however sampling those events statistically via BD simulation is at best inefficient and in some cases not possible.

In a human body, a relatively lower elongational flow rate seems to be ubiquitous at various positions throughout the vasculature, e.g., at the sites of stenosis, vasoconstriction, and in microvascular networks. Considering a very coarse estimate, the single-pass transit time of vWF molecules through those locations is likely in the range of milliseconds or less. While the probability of unraveling on a milliseconds timescale may be very low at lower elongational flow rate(22), periodic blood flow combined with the statistical nature of unraveling means that this probability must be considered many times during a typical vWF lifetime. Put differently, vWF protein total exposure time to such intermittent extensional flow can be on the order of minutes to an hour. This intermittent extensional flow exposure means the probability for pathological unraveling must be considered over longer timescales than are associated with single-pass events.

To analyze the probability of globular-unraveled transition of vWF multimers at relatively lower extensional rate, we have applied an enhanced sampling technique, i.e., the Weighted Ensemble method, in conjunction with Brownian dynamics (i.e., WEBD). This paper reports transition probability from globular to unraveled state per unit time computed from WEBD simulation for varied lower elongational rates. Further, we estimate the minimum extensional rate required to activate the healthy size vWF multimer for thrombus formation and cleavage by ADAMTS13. Our simulation prediction of threshold extensional rate is in good agreement with the experimental values(25).

## Methodology

### vWF Multimer Model

In simulations presented here, vWF monomers are modeled via a bead-spring description, as shown in Fig. 2; this model was presented previously but is briefly summarized(22). Two beads, connected by a parallel system of two springs, represent a monomer; the spring system consists of a finitely extensible non-linear elastic (FENE) spring and a relatively much stiffer harmonic spring(15). At sufficiently large applied tensile force, the harmonic spring is susceptible to a probabilistic fracture associated with the A2 domain initial unfolding event; for details, see Ref.(22). This model has been previously shown to accurately represent the complex mechanical response to pulling exhibited by A2 domains in the experiment. Each bead represents a collection of all domains on either side of the A2 domain. In this study, the focus is on the unraveling of vWF multimers at relatively lower extension rate flows, where such events in typical BD simulations are considered rare. Because of this, all A2-domains remain in a fully folded state such that the A2-domain mechanical response is dictated by the harmonic spring. Dimerization at the CT/CK terminus and multimerization at the D’D3 terminus are both modeled by connecting monomers with a stiff harmonic spring(26).

**Figure 2:**
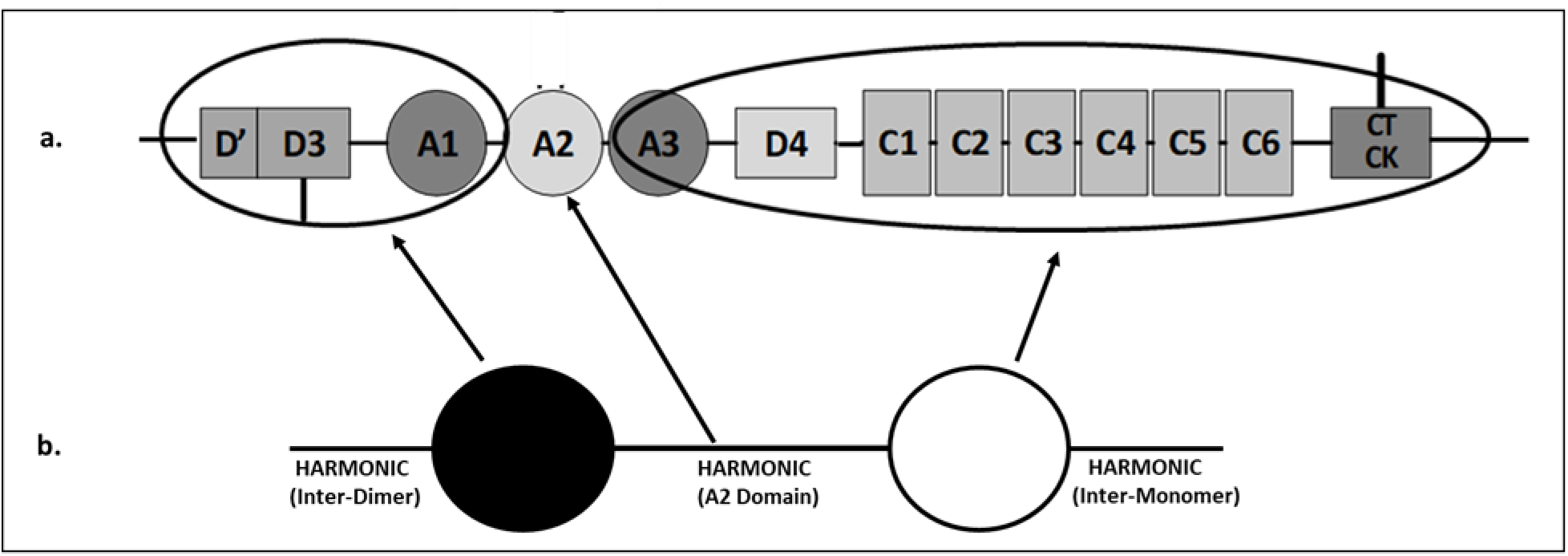
(a) Schematic illustration of vWF’s domains. (b) The vWF monomer model containing two rigid beads connected by a harmonic spring and arrow indicating what they represent in the model (see text).

### Brownian Dynamic (BD) Simulation

The enhanced sampling technique presented here was employed in combination with more standard Brownian dynamics (BD) simulations to model vWF protein or multimer, the response in varying strain rate elongation flow conditions. Details of the BD simulations are presented first.

In BD simulations, the dynamics of the i^th^ bead position **r_i_** is given by overdamped Langevin equation(27):

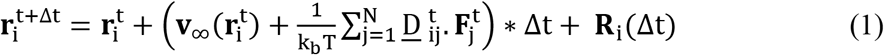

where, 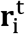 and 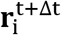 are the position of i^th^ bead at timestep t and t+Δt, respectively. **v_∞_(r)** is the undisturbed solvent flow profile and 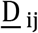 is the diffusion tensor; k_b_ is Boltzmann’s constant and T is the temperature. **R_i_(Δt)** is a random displacement with Gaussian distribution function whose average is 0 and variance-covariance is 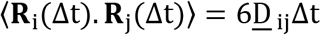. For elongational flow the undisturbed flow profile is 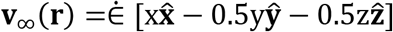, where 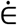 is the strain rate. x, y, z are the coordinates in x, y, z direction and 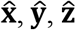 are the corresponding unit vectors. Hydrodynamic interaction among beads are manifested in 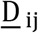 (diffusion tensor), which is given by the RPY approximation(28, 29):

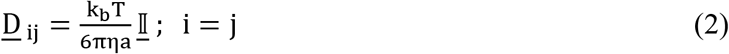

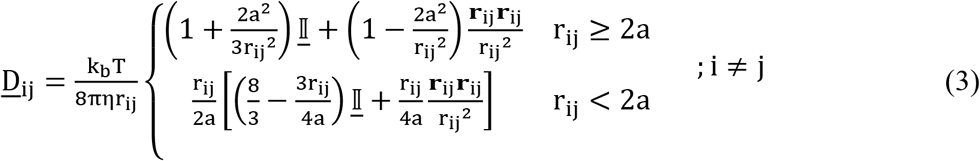

where i, j denote beads, **r_ij_ = r_j_ − r_i_**, 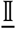 is the identity matrix and a is the bead radius.

**F_j_** in Eq. 1 represents the force acting on the j^th^ bead. **F_j_** is the summation of spring forces **F^s^** applied by adjacent beads and Lennard-Jones forces **F^LJ^** exerted by all the beads of a vWF multimer.

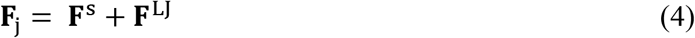

Beads are connected by two types of harmonic spring. One represents the A2 domain 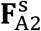 and the second is to model the disulfide bond between monomers 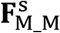.

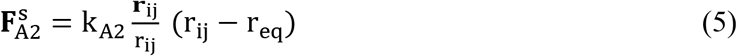

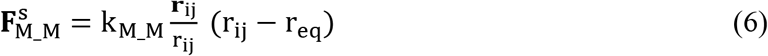

Experimental data for the force-extension correlation of folded A2-domain was used to compute k_A2_ = 5 pN/nm (15). To model the bond between the monomer as strong inter-disulfide bonds, the spring constant k_M_M_ is taken as k_M_M_ = 200k_b_T/a^2^. Both the harmonic spring oscillates around an equilibrium length r_eq_ = (2a + 1nm), which was elected to achieve an appropriate dimer length(30).

A modified 12-6 Lennard-Jones interaction is used for 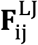:

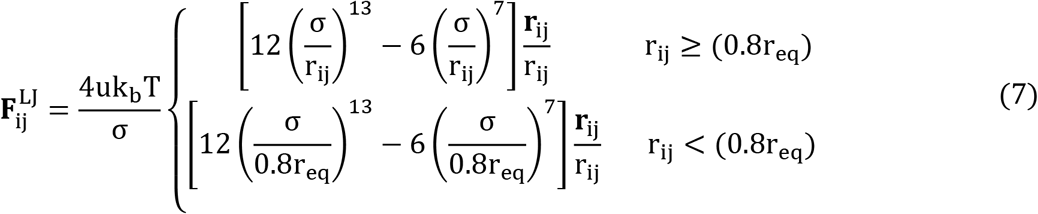

where u and σ are the energy and length parameter, respectively. The length parameter is 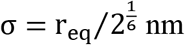. In quiescent or low flow rate conditions, vWF proteins exhibit highly compact, i.e., collapsed, conformations; this is achieved in the model by using energy parameter u = 1.0(19). The model of vWF has been parameterized on the monomer length scale, so the bead radius is taken as 15 nm(30), and its drag coefficient is determined utilizing Stokes drag law. The drag coefficient of the bead is ζ = 6πηa, where η is the solvent viscosity. We have selected solvent viscosity as η = 0.001pNμs/nm^2^, which is the viscosity of water, and the temperature is taken as T = 300K. The multimer length studied here is 80 monomers (i.e., 160 beads), which is near the upper end of the functional size range of vWF multimers found in blood plasma(7). Quantities are made dimensionless by rescaling lengths 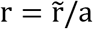 by the bead radius a, energies 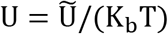 by thermal energy, and time 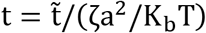 by characteristic diffusion time τ = ζa^2^/K_b_T.

### Weighted Ensemble Brownian Dynamics (WEBD)

The transition from globular to unraveled conformation at relatively lower elongation rates occurs on time scales potentially inaccessible to standard Brownian Dynamic (BD) simulations. As discussed in the Introduction, cases exist where the time scales are accessible but still relatively large; in such situations, BD simulations are inefficient for sampling elongation transitions. Hence, to examine the probability of unraveling at a low strain rate, we employed a weighted ensemble method, in combination with Brownian dynamics (WEBD). The idea of the WE method was presented by Huber and Kim as an alternative method for the simulation of protein-association reaction in which reaction events rarely occurred(31). Thereafter, a weighted ensemble strategy was used with varied stochastic dynamics to study various biological transition events that occur at timescales that are not feasible to conventional brute-force simulation methods(32–35). The reason for this shortcoming is the high energy barrier between stable/metastable states, as shown schematically in Fig. 3. The major advantage of WE method over standard stochastic dynamic simulation in the study of long-timescale events is that the computer resources in WE method are used uniformly over the entire configuration space independently of the free energy landscape. While in conventional simulation methods, most of the computational resources are utilized in the region of configuration space with low free energy. Hence, WE method is able to sample the region of high free energy, which is crucial for the study of transition events that are rare or may never be possible in conventional simulations. The detailed explanation of the WE method for the system studied here is given below.

**Figure 3:**
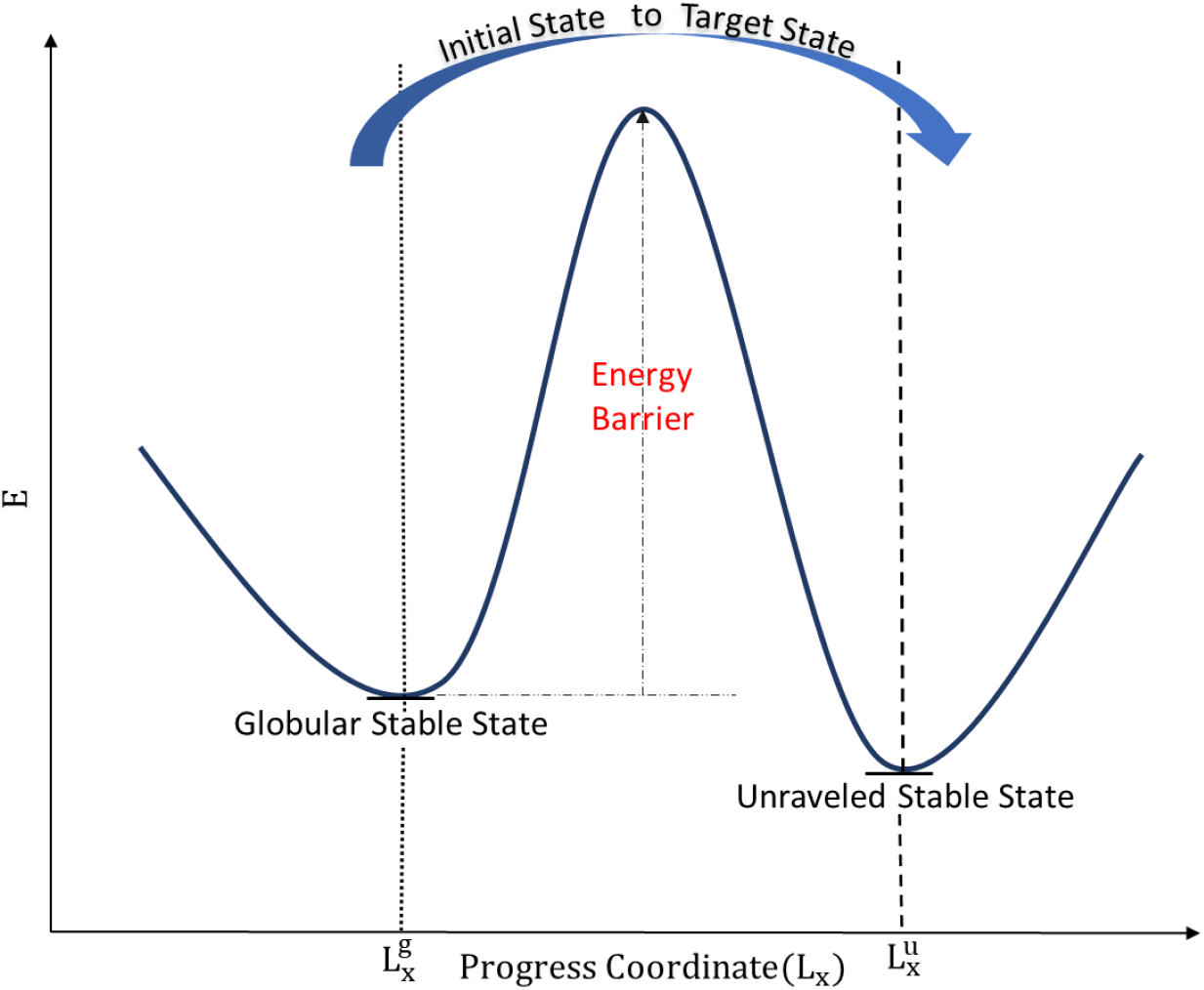
Blue solid line is for an energy profile as a function of progress coordinate for a vWF protein in elongational flows. In schematic example shown, the strain rate is considered sufficient high to make the unraveled state lower in energy than the globular state. Here, polymer extension in the flow direction (L_x_) is the progress coordinate. The initial state is defined as 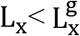, and target state as 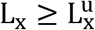.

The primary step in WEBD is to divide the configuration space into bins along the progress coordinate of the conformational transition event(31). We defined polymer extension (i.e., projected length of polymer) in the flow direction (L_x_) as the progress coordinate. Further, the initial and end (target) state were defined on the progress coordinate. Here, we study the transition event from one stable state (globular state) to another stable state (unraveled state). For elongation rates at which the unraveled state is a minimum on the energy surface, LU is well defined for that state; this is always true for the globular state because globule size is determined by multimer length and LJ interaction parameters. Hence, the initial state is defined as polymer extension less than that of globular state, and the target state as polymer extension equal to or greater than that of the unraveled state, as shown in Fig. 3. Computing the extension of polymer in the elongational strain direction for globular 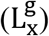 and unraveled stable states 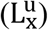 is described in the next section.

Besides defining the initial and target state, it is also essential to choose the number of bins and their placement along the progress coordinate before executing a WEBD simulation(31). For all extensional rates, the initial state is considered as the first bin while the target state as the last bin. To estimate the number of bins and their position between the first and last bin, we employed a Bin placement scheme proposed by Huber and Kim(31).

WEBD simulation begins with 20 non-interacting polymer chains that are initiated in the start state (i.e. 1^st^ Bin); their initial conformation was acquired from standard BD simulations of vWF multimers at a no-flow condition. The number of starting state samples used was equal to the user predetermined number of samples per occupied bin, M (see below). All these chains (i.e., initial state conformational samples) were assigned an equal probability such that the total is 1, as shown in Fig. 4.

**Figure 4:**
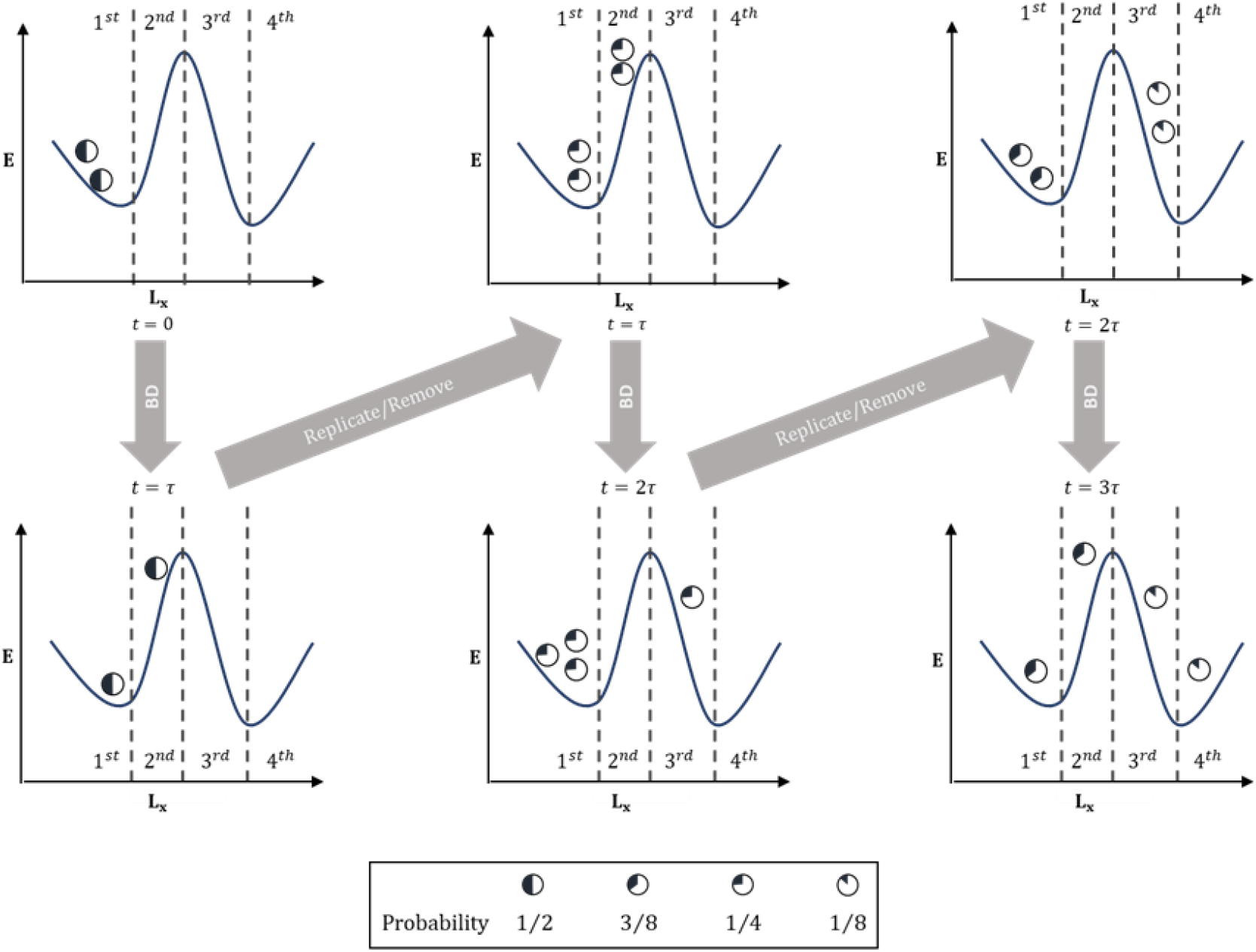
Schematic diagram of WEBD. In this example, configuration space is divided into four bins, and the number of polymer chains in each occupied bin is M = 2. Polymer chains are represented as circles and their assigned probability is proportional to the filled region of the circle. Note that the number of bins and the predetermined number of chains per occupied bin in this schematic are significantly less than the corresponding values used in WEBD simulations presented here.

These polymer chains undergo Brownian dynamics simulation for time increment τ = 10,000 BD steps. At each τ, the extension of each polymer chain in the flow direction is examined to determine the occupied bins. Thereafter, polymer chains are replicated or deleted to maintain the predetermined number of polymer chains (M) for each occupied bin, as shown in Fig. 4. In replication cases where the number of samples in that bin was less than 20, the replicated sample was chosen according to their existing probability; the same was true for all deletion cases. For newly created samples (via replication), a new seed for random number generation was used for the subsequent BD simulation. Following a replication/deletion step, conformational state probabilities were reassigned to maintain constant probability within a bin, which naturally conserves total probability across bins. The algorithm for replication/deletion of a polymer chain is exactly the same as the guidelines proposed in the original work of WE method for splitting/combining a particle(31).

To estimate transition rate from WEBD simulation, we have employed Hill’s relation, which states that the probability flux entering into the target state in a steady-state simulation is equal to the transition rate from the initial state to target state if the trajectories arriving at target state are fed back into the initial state(36). To apply Hill’s relation, a particular criterion must be used for polymer chains that reach the target state (i.e., the last bin). Throughout the simulation, polymer chains entering the target state are probabilistically reinitiated into the initial state by adding their probability to the initial state (i.e., the 1^*st*^ bin). The probabilities that are fed back into the initial state are distributed among its existing polymer chains. After some multiple τ steps, a steady state condition is reached in which probability per unit time of polymer chains that enter the target state fluctuates around a well-defined average value.

WEBD simulations are capable of computing transition rate for cases where standard BD simulation fails to do so; nonetheless, for WEBD simulation, the computer resources required to perform the study for lowest extensional rate examined here is significantly higher than those of highest rate explored here. On quantity that characterizes this, is the total number of samples generated during WEBD simulation; the total number of polymer chains for 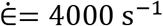 was 600, whereas for 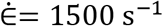 it was 4640. An alternative but related comparison is the total computational time where the total computational cost for 1500 s^−1^ was approximately 12 times than that of 4000 s^−1^.

### Globular and Unraveled Stable State’s Polymer Extension

In WEBD we have defined the initial and target state based on polymer extension (L_x_) in the flow direction of the globular 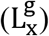 and unraveled state 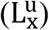, respectively. To assess the stability of the unraveled state as a function of elongation rate, BD simulations were performed wherein the starting state was a fully unraveled multimer with all constituent springs at their equilibrium values. vWF multimers were then subject to elongation flow of varying rates. Within the duration of the simulations (300ms), all chains remained unraveled for 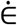 greater than or equal to 400 s^−1^; all samples modeled at 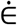 less than or equal to 300 s^−1^ collapsed to a globular conformation in the simulation duration. Hence, BD simulation suggests that unraveled configuration is unstable for an extensional rate less than or equal to 300 s^−1^ and at least metastable for an extensional rate greater than or equal to 400 s^−1^.

The extension of a polymer in an unraveled state depends on the elongation rate of the flow; Table 1 shows values computed from BD simulations described above. After a relatively short simulation, vWF multimers in a given flow rate condition converged on a steady-state ensemble-averaged polymer extension (Table 1). These simulations were also used to compute force profiles along the unraveled multimers, as presented in the next section. The globular state 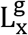 shows minimal dependence on 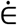; thus, for all strain rates examined here, the globular state’s polymer extension was taken to be the averaged extension of vWF multimers computed from quiescent flow simulations (i.e., 0.24μm).

**Table 1:** Ensemble Averaged polymer extension in the flow direction 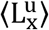 of unraveled state for varying extensional rate.

| Strain Rate($\text{s}^{-1}$) | $\langle L_x^u \rangle (\mu\text{m})$ |
| --- | --- |
| 400 | 4.10 |
| 500 | 4.34 |
| 1000 | 4.78 |
| 1500 | 4.95 |
| 2000 | 5.07 |
| 2500 | 5.18 |
| 3000 | 5.28 |
| 3500 | 5.38 |
| 4000 | 5.47 |

## Results and Discussion

### Threshold Extensional Rate for Activation of vWF

The goal here is to study the probability of multimer unraveling for healthy size vWF multimers at relatively lower elongational flow rates, at which unraveling is either difficult or not possible to observe in standard BD simulations. Further, for unraveled states, we wish to use information about the forces acting on A2 domains along the multimer to assess vWF activity (i.e., binding or scission).

In the absence of vascular injury, pathological vWF multimer unraveling may lead to thrombus formation or undesired protein scission. Experiments have shown that vWF binding to platelets at the A1 domain increases when the adjacent A2 domain is subject to some degree of tensile loading; furthermore, tensile loading at the domain scale is greater for unraveled multimers. Moreover, tensile loading on the A2 domain is also required for vWF scission because the A2 domain must exhibit significant domain unfolding to expose the Ter-Met scission site for interaction with ADAMTS13. Response to tensile loading of the A2 domain is characterized by an initially linear and relatively stiff mechanical response until a unique structure to the A2 domain undergoes an initial unfolding event. A2 domain mechanical response, including unfolding, has been previously explored via force spectroscopy experiments. Initial unfolding was observed to occur for applied forces between approximately 6 pN and 20 pN; the most probable initial unfolding force increased with increasing pull rate, but initial unfolding exhibited a probabilistic nature for all pull rates(15). After initial unfolding, subsequent A2 domain mechanical response was much more compliant such that, depending on the applied force at which initial unfolding occurred, the domain was observed to undergo significant extension in a constant force scenario.

As discussed in the preceding method section, BD simulations showed that the unraveled state is not a stable state for an extensional rate less than or equal to 300 s^−1^. For 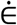 greater than or equal to 400 s^−1^, Fig. 5 shows plots of tensile force along an unraveled multimer; as has been observed previously, the force distributions for unraveled multimers in either elongation or shear flow exhibit a single peak at the center[16]. As Fig. 5 shows, the peak force acting on A2 domains exceeds the minimal unfolding force observed experimentally (i.e., 6 pN) for 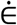 greater than 1000 s^−1^ and less than 1500 s^−1^. Experiments exploring A2 domain unfolding were executed over time on the order of minutes; the force abetted statistical nature of initial unfolding indicates that it may be observed for applied force less than 6 pN, but the rate of the reaction is relatively very low and continues to decrease for lower applied force(15). Results here indicate that elongation flow driven pathological vWF scission would not be observed for elongation flow rate equal to or below 1000 s^−1^. This prediction is in good agreement with recent experiments that explored vWF scission in varying extensional rate flow up to 1000 s^−1^; the authors observed no scission activity despite repeated exposure over durations of order 30 to 60 minutes(25). Furthermore, Fu et al., through microfluidic experiments gave evidence that the tensile force required to activate A1 domains for platelet adhesion is in a similar range to the tension required to unfold A2 domains(37). Hence, tensile force distributions shown in Fig. 5 also indicate that, for an extensional rate lower than 1500 s^−1^, the probability for binding of platelet to A1 domain will be negligible. Consequently, thrombosis formation for those elongational rates is not expected.

**Figure 5:**
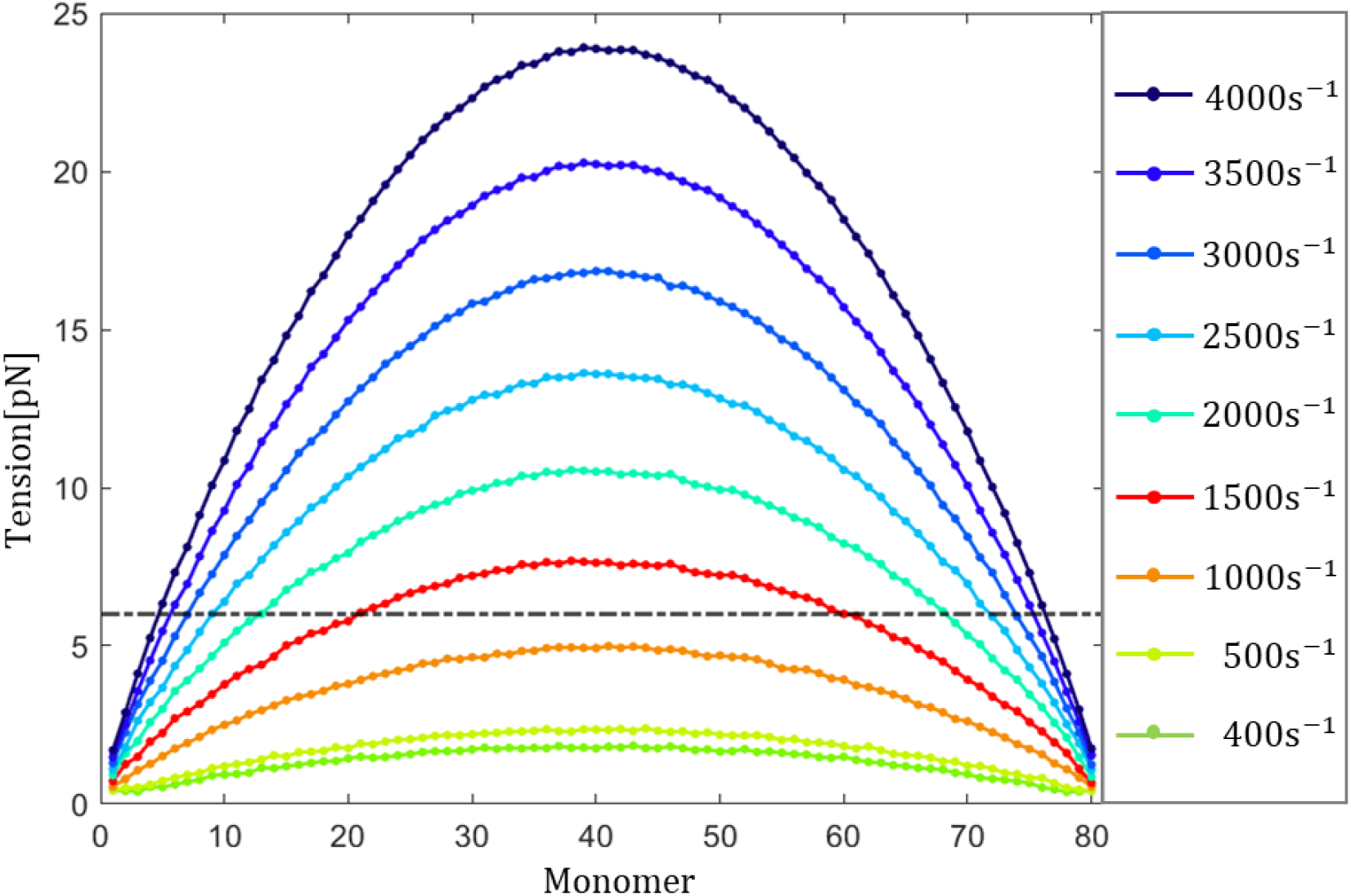
Ensemble averaged tensile force distribution along unraveled vWF multimers for varying extensional rate. The dashed-dot line shows the lowest applied force value for which A2-domain unfolding was observed in prior experiments.

Employing WEBD, we compute the probability of multimer unraveling at a relatively low elongational flow rate to estimate the risk of cleavage of healthy size vWF and thrombus formation in blood vessels. Tensile force along the multimer suggests that for 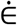 less than 1500 s^−1^, the probability of those risks is negligible. Hence, here we have examined the transition from globular to the unraveled state of vWF molecule only for elongational flow rates where thrombus formation and cleavage by ADAMTS13 is considered possible (i.e. 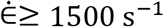).

### Transition Rates from a Globular to an Unraveled State

An overarching goal of this work is to predict transition rates for going from a globule to an unraveled state for vWF in elongation flow. A WEBD simulation must reach a steady-state condition before a transition rate between states can be computed. Steady-state is reached when all ensemble bins have time-independent probability; furthermore, the average probability flux entering the target state (i.e., the last bin) must be time-independent.

Figs. 6(a and b) illustrate the flux into the target state and the occupation probability of the 1^*st*^ bin as a function of time, respectively, as functions of time. After a relatively short simulation duration, both values fluctuate around well-defined average values. However, the target state probability flux exhibits a relatively high variance, particularly for the lowest rates explored here. The reason for such high fluctuations around the average value is the significant variation in first passage time (i.e., the inverse of target state probability flux(36, 38)) for the globular-unraveled transition(20); this variation increases with decreasing elongational flow rate. Hence, the precise way to confirm steady-state has been achieved in a WEBD simulation is to compare the distribution of observed probability flux into the target state computed for two different time ranges in the data set. The results of such an analysis are shown in Fig. 7 and, for all rates explored, a minimal difference exists between the distributions obtained over two different ranges in time. For the lowest rates explored, there appear to be some differences in the distribution of lower tails, but the average predicted for both distributions is the same. While Fig. 7 confirms that most of the simulation trajectory for each case shown in Fig. 6a is in a steady-state; the simulation durations shown were necessary to obtain acceptably small error bars for computed averages. When a WEBD simulation reaches the steady-state condition, the probability flux entering the target state is equal to the probability of a transition from a globular to an unraveled state in one second(36).

**Figure 6:**
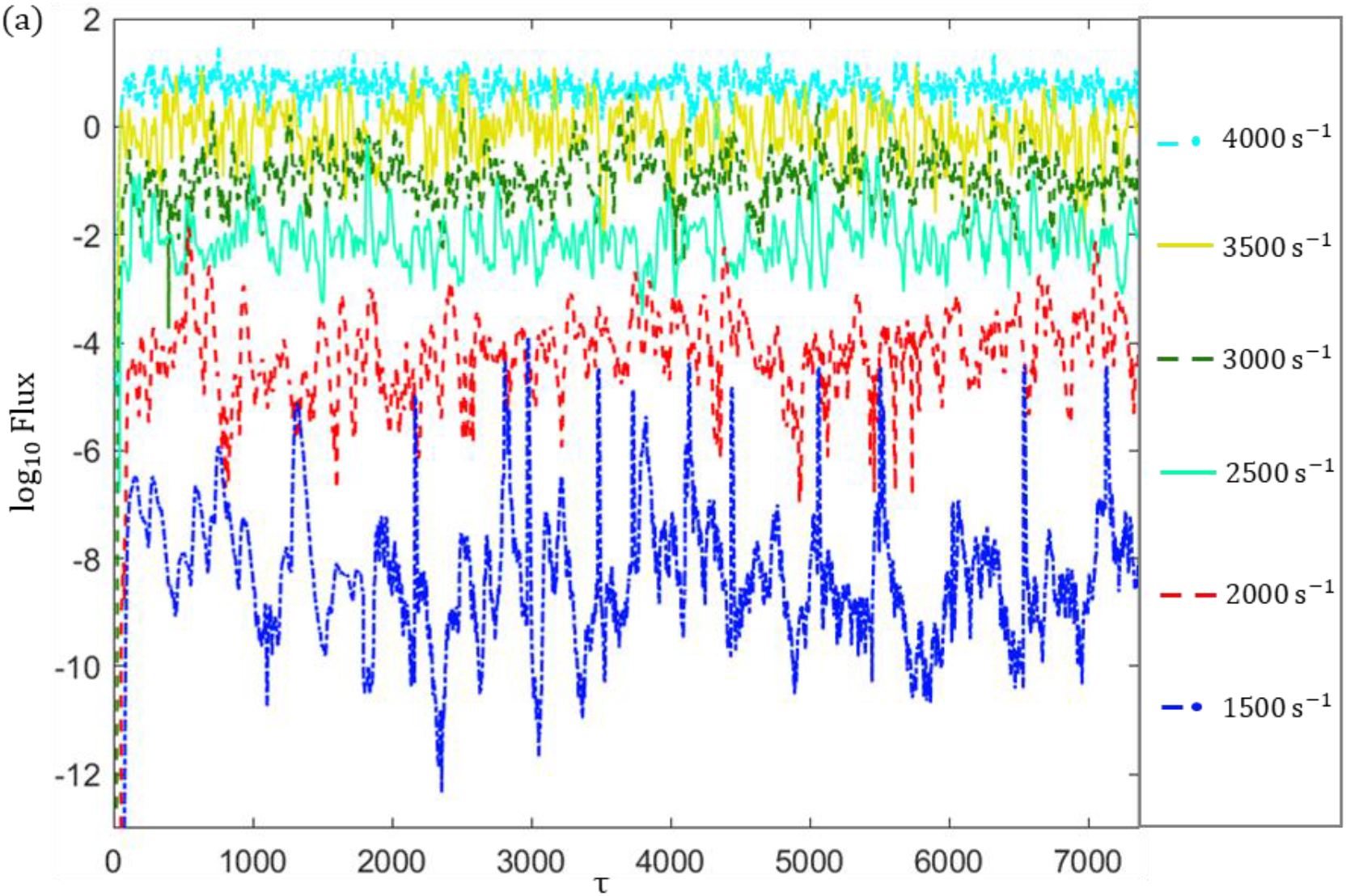
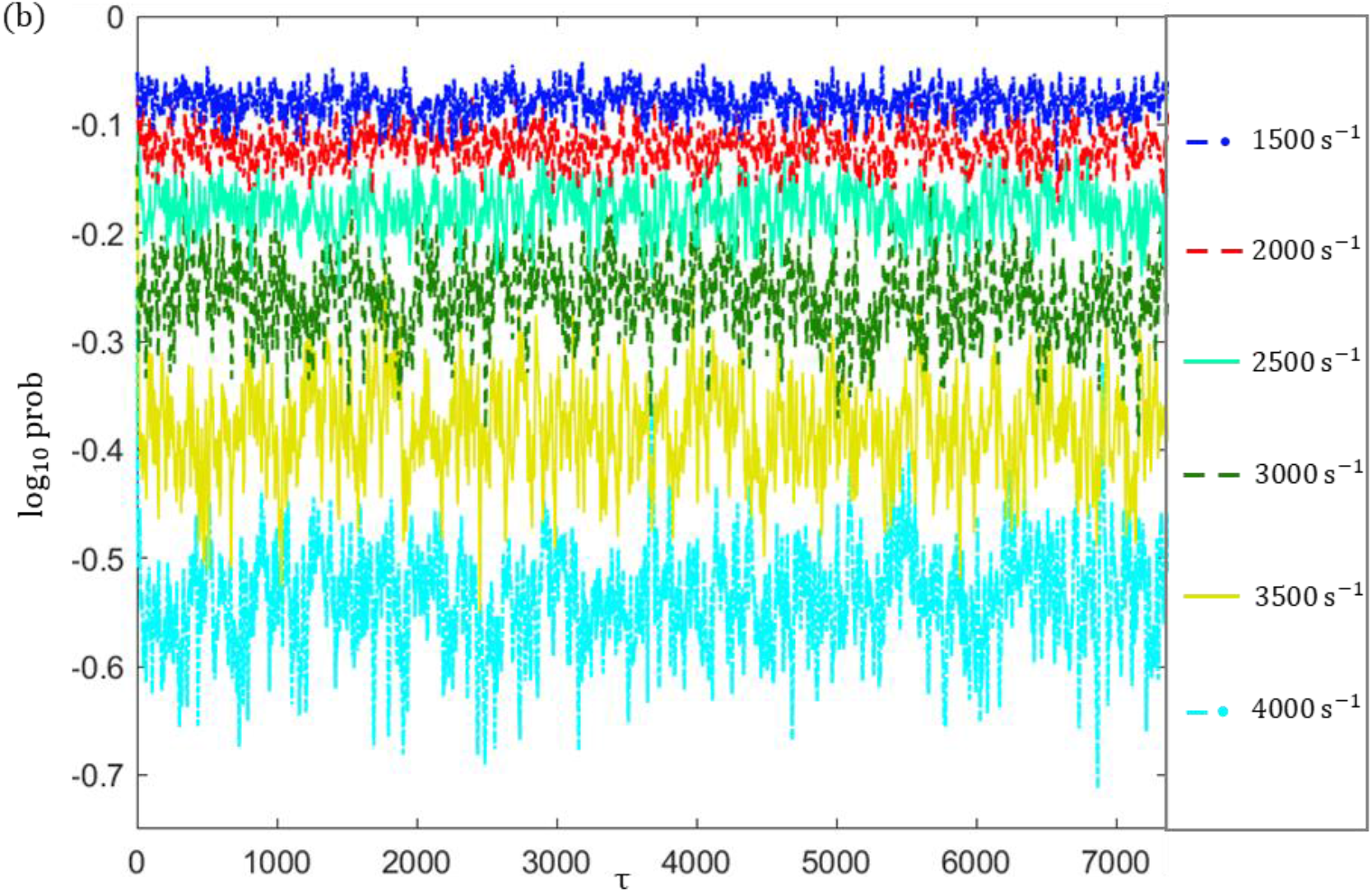
(a) Probability flux into the target state as a function of time for all extensional rate. (b) Probability of 1^*st*^ bin as a function of time. For all strain rates probability flux and probability are window-averaged over 10τ.

**Figure 7:**
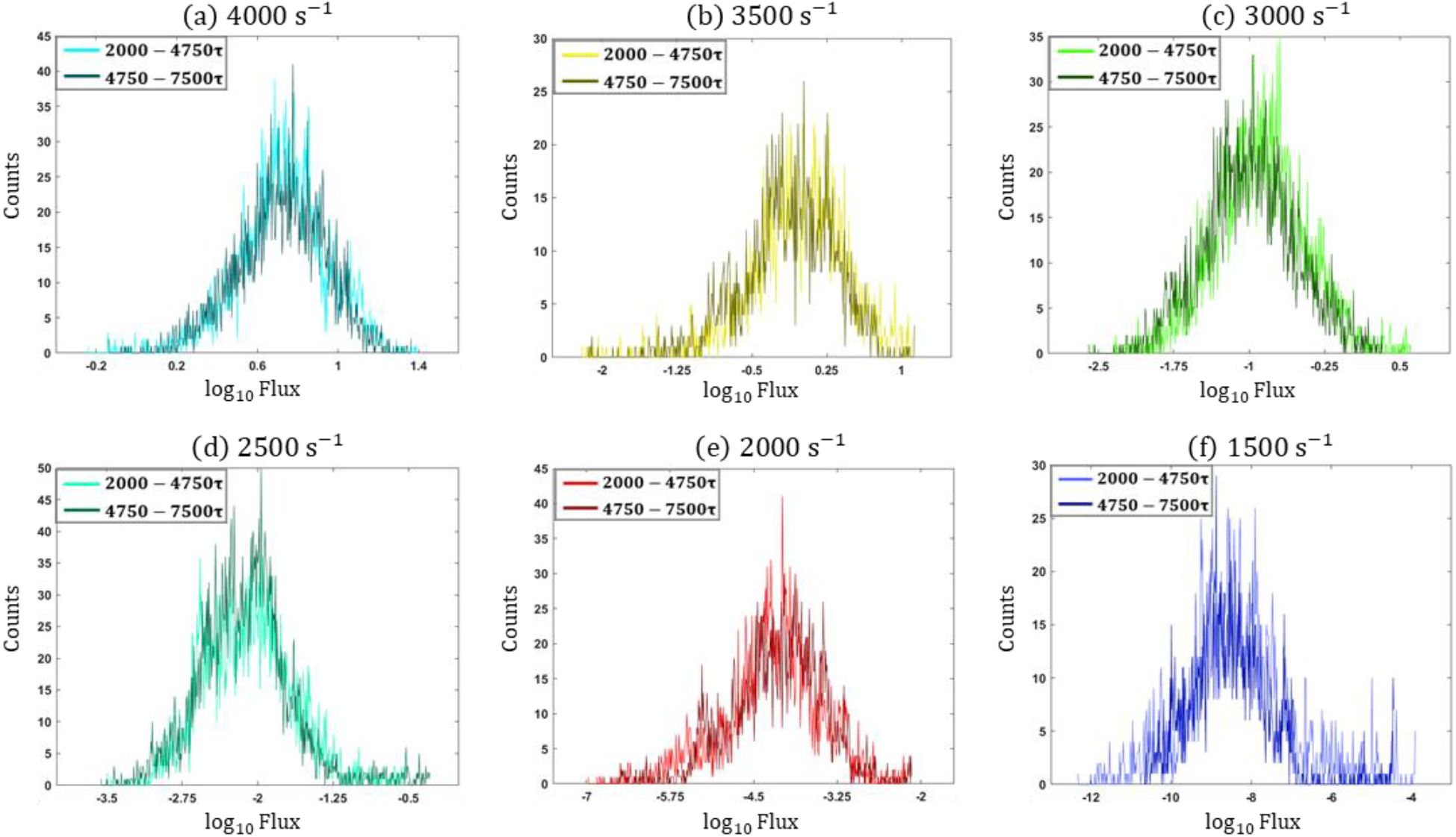
(a-e) Distribution of target state probability flux for time range [2000τ – 4750τ] and [4750τ — 7500τ]. All results shown here for probability flux are window-averaged over 10*τ*.

As Figs. 6(a) and 7 show, the target state probability flux decreases dramatically as the elongation flow rate decreases, in accord with the reduction in likelihood for a globular to unraveled transition being observed at a lower extension rate. Note that for the highest rates explored 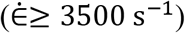, the log of the predicted probability flux is greater than zero, indicates a probability greater than unity. This should instead be interpreted that the mean first passage time for the globular to unraveled transition is less than 1 second at 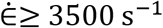. This is addressed further below when presenting average transition times. Figure 6(b) shows that the occupation probability for the first bin follows an inverse behavior as to what is exhibited for the target state probability flux shown in Fig. 6(a). This illustrates the strongly increasing probability of remaining in the globule state for a lower extensional rate of flow.

To cast this discussion in terms of time, Fig. 8 illustrates the distribution of transition rate (/sec) from a globular to an unraveled state. Transition events from a globular to an unraveled state can also be viewed as a form of instability, similar to material failure (in cases explored here, the load is caused by hydrodynamic forces). Failure rates in many processes have been observed to follow a lognormal distribution(39, 40), and Fig. 8 shows the same is true for the transition rate from a globular to an unraveled conformation. Figure 8(a,b) shows the probability distribution data for the log of the transition time along with normal fits (i.e., a lognormal behavior in the transition rate). Figure 8(c) bears this out further by comparing the observed cumulative distribution functions along with their fit curves.

**Figure 8:**
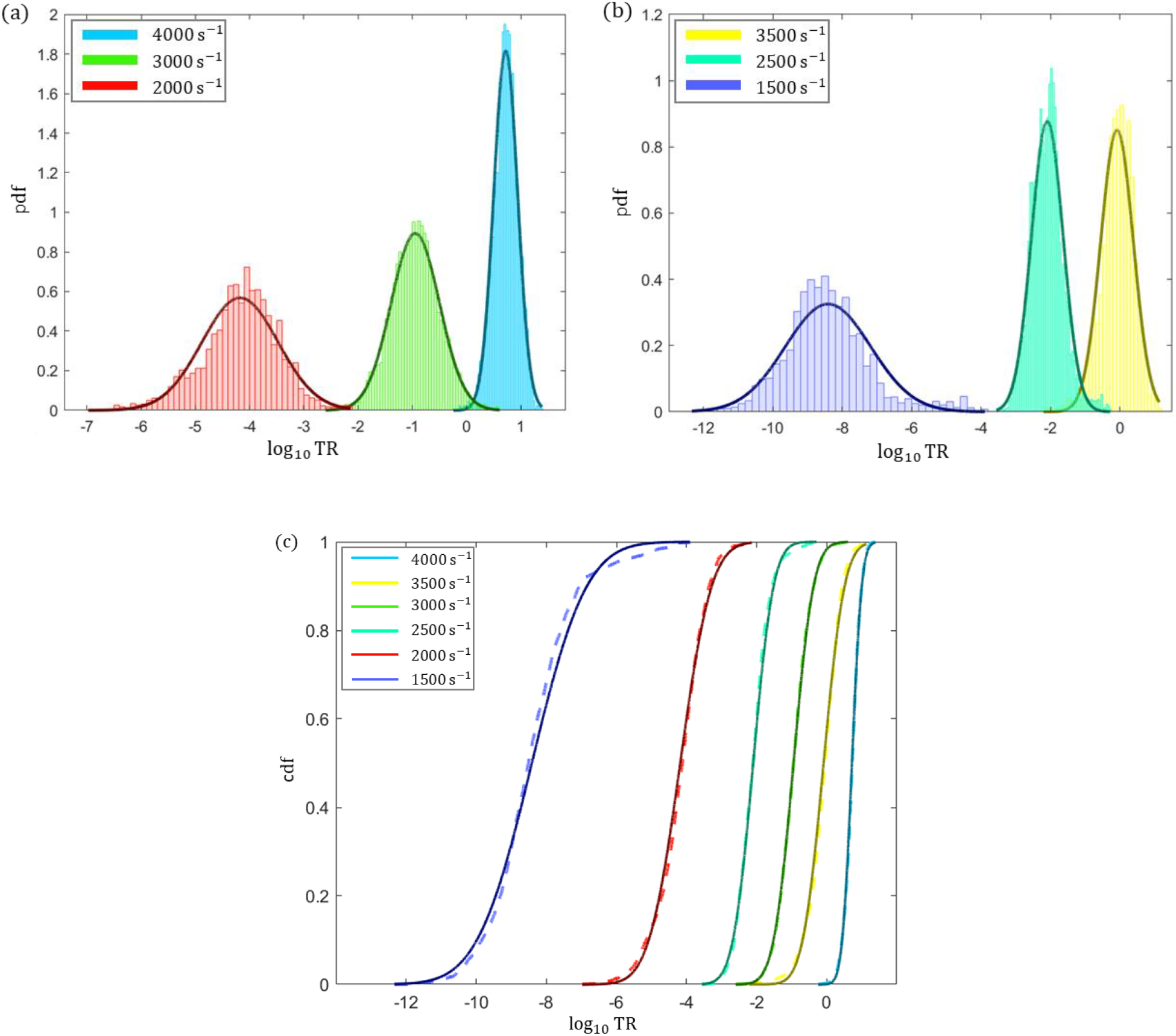
(a,b) Probability distribution function (pdf) of transition rate that can be well fitted with a normal distribution (solid line). (c) Comparison of cumulative distribution functions (cdf) of transition rate (dashed line) with the cdf of fitted normal distribution (solid line).

For all extensional rates explored, we computed the mean transition rate 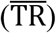 and results are shown in Fig. 9. For the globular-unraveled transition, the polymer chain has to overcome an energy barrier, as shown in Fig. 3. One way to interpret the influence of flow and the corresponding applied force on a vWF multimer is that there is a reduction in the energy barrier with increasing extensional rate, which causes the transition rate to increase. The highly non-linear dependence of transition rate on the strain rate suggests that elongation flow significantly influences the energy landscape of the globular-unraveled transition.

**Figure 9:**
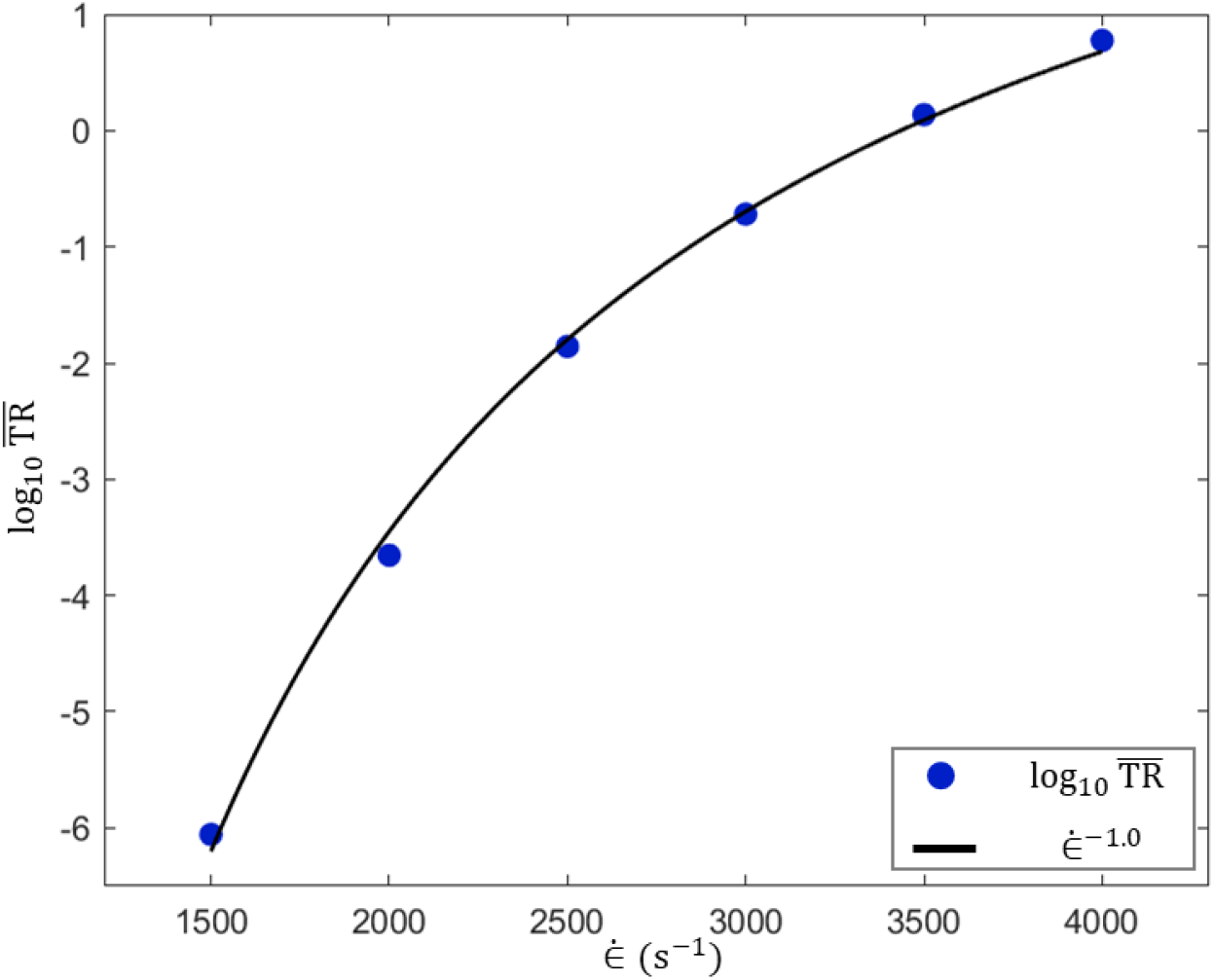
Mean transition rate 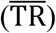 as a function of strain rate.

An additional way to interpret data obtained in the WEBD simulations is to compute the time for which the average probability of transition equals unity. In other words, in that amount of exposure time to a given rate extensional flow – on average – a vWF multimer would undergo unraveling. Those predicted times are shown in Table 2, and the highly non-linear dependence on the extensional flow strain rate is evident.

**Table 2:** Averaged exposure time required to unravel vWF multimer for varied extensional rate.

| Strain Rate($s^{-1}$) | Required Exposure time |
| --- | --- |
| 1500 | 315.8 hours |
| 2000 | 1.25 hours |
| 2500 | 1.20 minutes |
| 3000 | 5.4 seconds |
| 3500 | 0.73 seconds |
| 4000 | 0.16 seconds |

It is important, when interpreting the times reported in Table 2, to acknowledge that vWF molecule doesn’t need to be consistently exposed to the given strain rate for the reported duration for the transition to occur. The globular-unraveled transition is of course statistical in nature; a given vWF molecule may exhibit a transition in a significantly shorter period than reported in Table 2 – or it may take much longer. One must also consider the periodic nature of blood flow (i.e., blood circulates through the human body of order twice per minute) and the reported half-life of vWF proteins in human circulation (i.e., 8 to 12 hours(23)). It is also useful to consider the highly complex flow to which vWF proteins may be subjected as they circulate among much larger blood entities. For example, flow between two approaching red blood cells may expose a vWF protein to condition with a very elongation rate but for a very short duration. At a scale local to a given protein, flow conditions and rates can exhibit more significant variation – spatially and temporally – than might be derived from considering average blood flow through the entire blood vessel(41). Even at the vessel scale, flow near bifurcations(42) and narrowing blood vessels, particularly associated with stenosis or other partial blockages, can generate extensional flow with a high strain rate.

Table 2 reports that exposure to extensional flow with strain rate 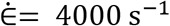 will likely drive unraveling in less than 160 ms; while such a flow rate may seem highly pathological, it may not be for very short duration events at the individual vWF protein scale. If such events occur with some regularity, they may lead to periodic vWF unraveling and associated pathology. Given the force data in Fig. 5, the unraveled state at 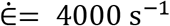 is highly activated for both platelet binding and ADAMTS13 scission. At 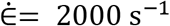, the time reported is much longer, but periodic exposure to such an extensional flow rate may be more frequent and persist for longer durations. The time indicated in Table 2 for 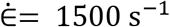 is of particular note because it is significantly larger than the reported vWF half-life. Data in Table 2 indicate that infrequent but regular exposure to extensional flow fields with strain rate approximately 1500 s^−1^to 2000 s^−1^ may represent a natural mechanism by which functional sized vWF proteins undergo scission and are recycled in the bloodstream. Further, periodic exposure to extensional flow with 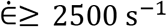 may represent mechanisms of pathology.

## Conclusions

We employed WEBD simulations to study the transition event of vWF multimer from globular to unraveled state at relatively lower extensional rate, such that observing transitions is computationally constrained – or even not possible – in standard Brownian dynamics simulations. For each extensional rate studied, the distribution of the transition rate follows a lognormal distribution. Moreover, the mean transition rate 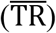 increases with the increment in the strain rate, according to the scaling law log_10_ 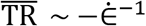. For all strain rates studied here, we computed the average total time required for vWF multimer to be exposed to either continuous or intermittent extensional flow to induce unraveling, which in turn can assist in estimating the likelihood of unraveling for a healthy size vWF multimer during its entire lifetime. Furthermore, from the tensile force profile of unraveled multimer, the threshold extensional rate required to activate functional size vWF for thrombus formation and cleavage was estimated. Collectively, results obtained here indicate that pathological scission will not occur for extensional rate below approx. 2000 s^−1^, which is in agreement with data from a recent experiment(25). The results presented here could provide important interpretation for the development of treatment for numerous diseases related to blood clotting and excessive cleavage of functional size vWF multimers caused by pathological flow condition.

## Author Contributions

All authors contributed to the design of this study. S.K. performed WEBD and BD simulations and data analysis. E.W., A.O., X.C., and X.F.Z. critically revised the manuscript and offered experimental/computational expertise.

## Acknowledgments

Computer resources were provided by the Extreme Science and Engineering Discovery Environment (XSEDE), which is supported by the National Science Foundation, Grant No. ACI-1548562(43). Moreover, support from the Lehigh sol cluster is also acknowledged.

